# Sperm cryopreservation impacts the early development of equine embryos by downregulating specific transcription factors

**DOI:** 10.1101/2021.05.12.443855

**Authors:** José M Ortiz-Rodríguez, Francisco E. Martín-Cano FE, Gemma Gaitskell-Phillips, Álvarez Barrientos A, Heriberto Rodríguez-Martínez, Maria C. Gil, C Ortega-Ferrusola, Fernando J. Peña

## Abstract

Equine embryos were obtained by insemination with either fresh or frozen-thawed spermatozoa at 8, 10 and 12 h post spontaneous ovulation, maintaining the pairs mare-stallion for the type of semen used. Next generation sequencing (NGS) was performed in all embryos and bioinformatic and enrichment analysis performed on the 21,058 identified transcripts. A total of 165 transcripts were downregulated in embryos obtained with cryopreserved spermatozoa respect embryos resulting from an insemination with fresh spermatozoa (p=0.021, q=0.1). The enrichment analysis using human orthologs using g:profiler on the downregulated transcripts marked an enrichment in transcription factors (TFs) in mRNAs downregulated in embryos obtained after insemination with cryopreserved spermatozoa. The 12 mRNAs (discriminant variables) most significantly downregulated in these embryos included among others, the *chromatin-remodeling ATPase INO80, Lipase maturation factor 1 LMF1, the mitochondrial mRNA pseudouridine synthase RPUSD3, LIM and cysteine-rich domains protein 1, LMCD1*. Sperm cryopreservation also caused a significant impact on the embryos at 8 to 10 days of development, but especially in the transition from 10 to 12 days. Overall, our findings provide strong evidence that insemination with cryopreserved spermatozoa poses a major impact in embryo development that may compromise its growth and viability, probably due to modifications in sperm proteins induced by cryopreservation. Identification of specific factors in the spermatozoa causing these changes may improve cryopreservation.

## INTRODUCTION

Cryopreservation is widely used due to their importance for the international commerce of equine semen of superior sires, and as a safety measure to avoid loses of genetic material in case of accidental death of a stallion. However, this technology still is suboptimal and numerous drawbacks persist (Pena et al. 2011). The fertility obtained with this technology is overall considered reduced compared with fresh-extended spermatozoa, and one of the causes is related to increased embryo loss when frozen semen is used (Panzani et al. 2014; Ortiz-Rodriguez et al. 2019b). In addition, delayed embryo development is associated with insemination of cryopreserved spermatozoa (Stout 2006). A growing body of scientific evidence indicates that the spermatozoa have important regulatory roles in the early embryo development (Jodar et al. 2020). These findings underpin the importance of potentially damaged sperm factors controlling early embryo development (Castillo et al. 2018). Processing the ejaculate for cryopreservation implies the removal of seminal plasma, extension in media containing cryoprotectants cooling, freezing, storage in liquid nitrogen, and thawing. All these procedures impact the spermatozoa; numerous mechanisms explaining cryodamage have been described, including toxicity of cryoprotectants (Macias Garcia et al. 2012), osmotic shock at freezing, and specially at thawing damaging membranes and the mitochondria (Garcia et al. 2012; Pena et al. 2015). This mitochondrial damage causes redox imbalance and accelerates different forms of sperm death, including necrotic, apoptotic (Aitken and Koppers 2011; Aitken and Baker 2013) and probably ferroptotic sperm mortality (Ortiz-Rodriguez et al. 2020). In addition to protein degradation during cryopreservation, redox deregulation may induce oxidative damage to proteins causing notable modifications of the sperm proteome (Bogle et al. 2017; Martin-Cano et al. 2020a; Gaitskell-Phillips et al. 2021b). All these modifications may affect proteins participating in the regulation and or survival of the early embryo (Castillo et al. 2018). Thus, we tested the hypothesis that early embryos, product of insemination with cryopreserved spermatozoa display detectable transcriptomic alterations that could jeopardize their development, taking in account that RNA seq is considered a relevant tool to evaluate embryo competence (Groff et al. 2019).

## MATERIAL AND METHODS

### Embryo collection and experimental design

Animals were maintained according to European regulations, and all experimental procedures were reviewed and approved by the Ethical committee of the University of Extremadura, Cáceres, Spain. Mares were treated with a prostaglandin analogue to shorten the luteal phase and were monitored daily for follicular development, degree of uterine edema and cervical tone using transrectal ultrasonography (US). When a follicle at least 35 x35 mm was detected in absence of luteal tissue, with good uterine edema and low cervical tone, mares received 2.500 IU of hCG. Follicular development was thereafter closely monitored by US and mares were inseminated immediately once ovulation was detected either with fresh sperm or frozen-thawed sperm from the same stallion (two mares were paired per stallion, one for each type of sperm used). A total of 18 conceptuses were obtained by uterine lavage at 8, 10, or 12 days post ovulation; embryos were snap frozen in LN_2_ and stored at −80°C until analysis.

### Isolation of RNA

Total RNA was isolated from the embryos using the kit PicoPure™ RNA Isolation Kit (Catalog number:KIT0204, Thermofisher) following manufacturer instructions. RNA concentration and quality were assessed by automatic electrophoresis using 2100 Bioanalyzer (Agilent Technologies, Santa Clara, CA, USA).

### RNA-seq analysis

Libraries were built for RNA-seq analysis in an IonTorrent S5/XL sequencer (Thermo Fisher Scientific, Waltham, MA USA). The raw reads were aligned to a horse transcriptome generated using ENSEMBL (Equ Cab 3 version) in Torrent server with the proprietary ThermoFisher algorithms. Then, BAM files were imported into Qlucore Omics Explorer ver 3.7 (https://www.qlucore.com) for analysis.

### Bioinformatic Analysis

#### Variance filtering and PCA

Transcripts were normalized to TPM, and then aligned data were normalized and log_2_ transformed using Qlucore Omics Explorer (https://qlucore.com). Principal Component Analysis (PCA) was used to visualize the data set in a three-dimensional space, after filtering out variables with low overall variance to reduce the impact of noise and centering and scaling the remaining variables to zero mean and unit variance. The projection score (Fontes and Soneson 2011) was used to determine the optimal filtering threshold.

#### Identifying discriminating variables

Qlucore Omics Explorer (https://qlucore.com) was used to identify the discriminating variables with significant differences between transcripts in embryos resulting from inseminations with fresh or cryopreserved spermatozoa, and in embryos of different ages also either obtained with fresh or frozen thawed spermatozoa. The identification of significantly different variables between the different subgroups of embryos was performed by fitting a linear model for each variable semen used in the insemination (fresh or frozen thawed) and age of the embryo. P-values were adjusted for multiple testing using the Benjamini-Hochberg method (Viskoper et al. 1989; Tamhane et al. 1996) and variables with adjusted p-values equal or below 0.1 were considered significant.

### Gene Ontology and pathway analysis

PANTHER (http://www.pantherdb.org/pathway/pathwayList.jsp) and KEGG pathway (https://www.genome.jp/kegg/) (Ogata et al. 1999; Altermann and Klaenhammer 2005; Du et al. 2014; Mi et al. 2019) analysis was used to identify biological pathways likely to be active in the proteins enriched in each group. Reactome (https://reactome.org) (Ogata et al. 1999; Altermann and Klaenhammer 2005; Du et al. 2014; Mi et al. 2019) analysis were used to identify biological pathways likely to be active in the transcripts enriched in each group, g:Profiler was also used to perform an enrichment analysis (Raudvere et al. 2019). Due to the increased depth of the human proteome in terms of annotation, the equine annotations were transformed to their human orthologs using g:Profiler (https://biit.cs.ut.ee/gprofiler/orth) and a pathway enrichment analysis and visualization was performed again using g:Profiler and Cytoscape analysis using Reactome (https://reactome.org).

### Network analysis

Cytoscape (https://cytoscape.org) plug in ClueGo was used to identify functionally grouped gene ontology terms in equine seminal plasma as described in (Bindea et al. 2009; Mlecnik et al. 2018). STRING (https://version-10-5.string-db.org) was used to identify potential functional partners of specific proteins.

## RESULTS

### Overall impact of cryopreserved spermatozoa on the transcriptome of equine embryos

A total of 21058 transcripts were identified. In a first step we constructed volcano plots to have a general overview of the impact of the type of semen (fresh or cryopreserved) on the transcriptome of the equine embryos (Figure 1 B). Then we compared the overall change induced by insemination with frozen and thawed spermatozoa on the embryonic transcriptome; we found that 165 transcripts were downregulated in embryos obtained with cryopreserved spermatozoa respect embryos resulting from an insemination with fresh spermatozoa (Figure 1AC; p=0.021, q=0.1). We performed enrichment analysis, using human orthologs, on the transcripts downregulated in embryos using g profiler, and significant enrichment was detected in different gene ontology (GO) terms; particularly notable was the enrichment in transcription factors (TFs) in transcripts downregulated in embryos obtained after insemination with cryopreserved spermatozoa. A total of 84 TFs were identified (Figs 2 and supplementary figure 1), including as most significantly enriched the TF NF-1 (p=1.040×10^−14^), TF KLF13 (p=2.734×10^−14^), TF CPBP (p=1.450× 10^−12^), TF BTEB3 (p= 1.139×10^−11^), TF TCF7L1 (p=1.398× 10^−11^) and TF KLF3 (p= 5.8× 10^−11^).

**Figure 1.-.**
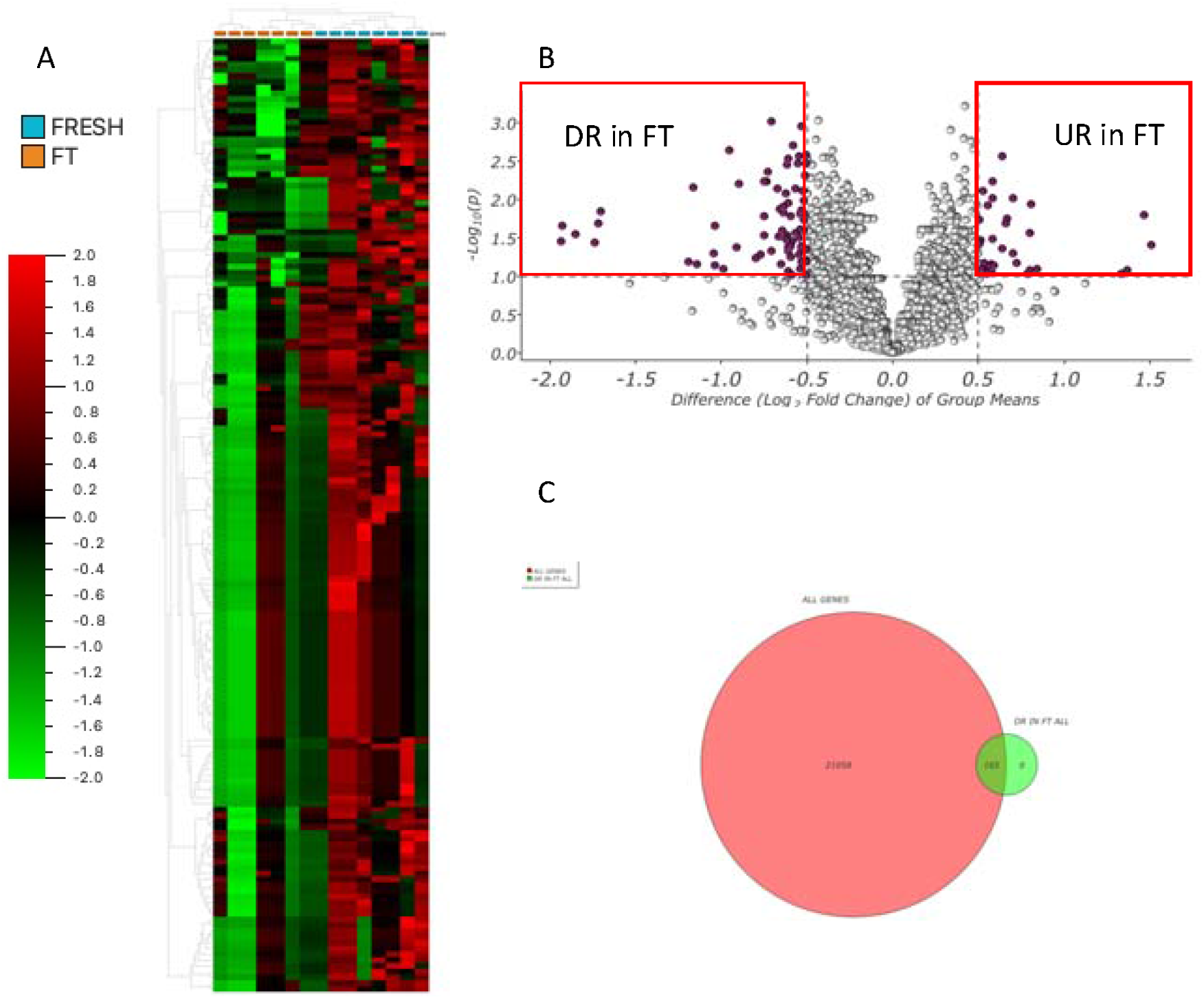
Changes in the embryo transcriptome depending of the type of semen used, fresh or cryopreserved. A) Heat map showing different expression of transcripts in embryos produced with fresh semen (right part), and those obtained with cryopreserved sperm (left part). In B, a Volcano plot showing transcripts differentially expressed in embryos obtained with fresh or cryopreserved spermatozoa. C) Venn diagram showing transcripts downregulated in embryos obtained with cryopreserved spermatozoa.

**Figure 2.-.**
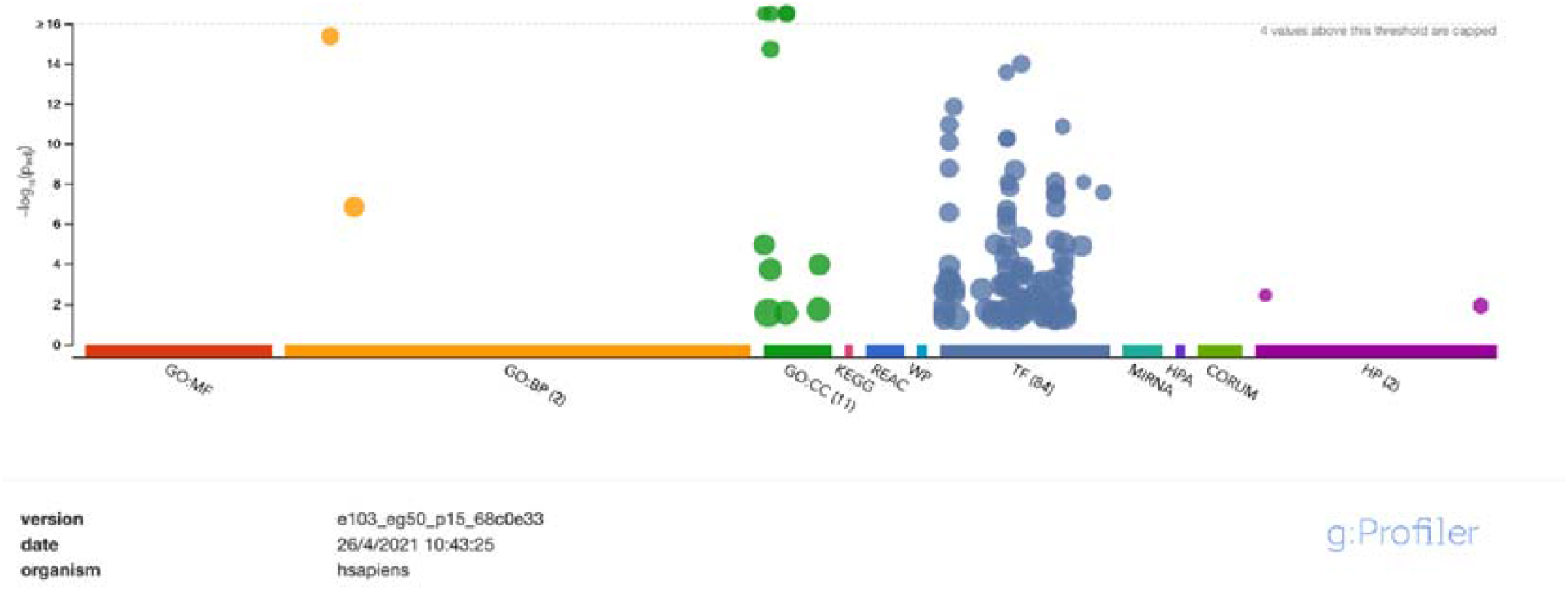
g:GOST Manhattan plot showing terms enriched in downregulated transcripts in embryos obtained with cryopreserved spermatozoa (human orthologs were used due to the deeper annotation of the human genome). Interestingly, 84 transcription factors appeared downregulated. Detailed information appears in supplementary figures 1 and 2

### Network visualization analysis

The visualization of integration networks using the Cytoscape app Clue go revealed annotations like regulation of nuclear division, mitotic DNA damage checkpoint and positive regulation of molecular mediator of immune response, and interleukine 4 and 13 signalling were significantly affected by sperm cryopreservation (Fig 3)

**Figure 3.-.**
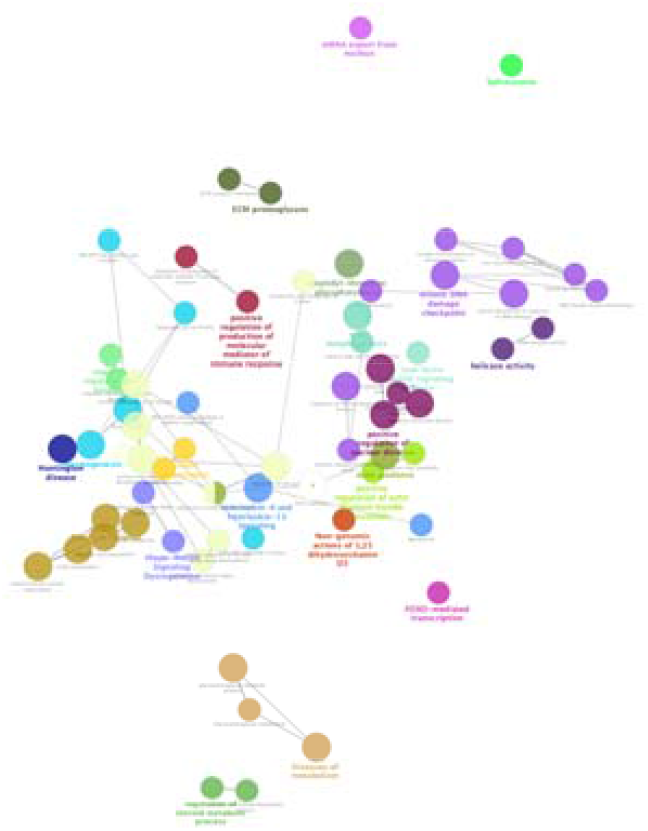
ClueGo network analysis of proteins upregulated in embryos obtained with fresh semen compared wth respect to those in embryos obtained with cryopreserved spermatozoa. To reduce the redundancy of GO terms, the fusion option was selected. GO/KEGG/REACTOME pathways functionally grouped networks with terms are indicated as nodes (Benjamini-Hockberg P value <0.01) and a minimum of 5 genes per group, linked by their kappa score level >0.4 where only the label of the most significant term per group is shown.

### Identification of discriminant variables between fresh and cryopreserved spermatozoa derived embryos

To reduce the number of variables, we applied stricter statistical criteria to reduce the number of variables differing between both conditions. We identified 12 discriminant variables significantly downregulated in embryos obtained with cryopreserved spermatozoa including among others, the *chromatin-remodeling ATPase INO80, Lipase maturation factor 1 LMF1, the mitochondrial mRNA pseudouridine synthase RPUSD3, LIM and cysteine-rich domains protein 1, LMCD1* (Figure 4).

**Figure 4.-.**
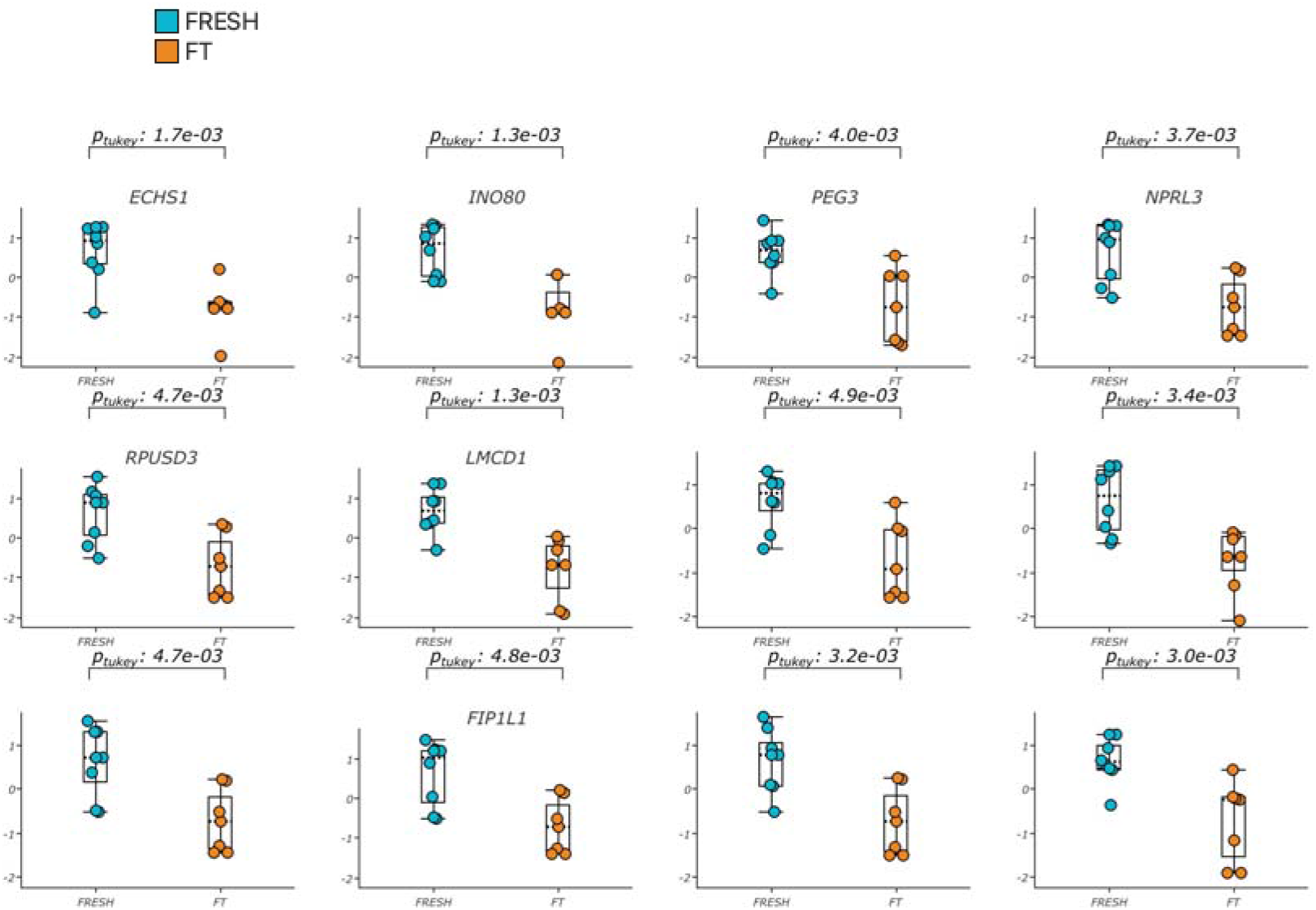
Differences in the amounts of specific transcripts in embryos obtained with inseminations with fresh or cryopreserved spermatozoa.

### Impact of cryopreservation on the development of equine embryos from 8 to 10 days post ovulation

We investigated if the use of frozen thawed spermatozoa impacted embryo development in two developmental stages 8 to 10 days and 10 to 12 days after ovulation. We identified numerous genes differentially expressed in both stages. On the first stage studied, transition 8 to 10 days, 202 upregulated expressed transcripts were detected (Fig 5A; p=0.04 q=0.1) in embryos obtained after insemination with fresh semen, while in embryos obtained after insemination with frozen thawed semen only 39 transcripts were upregulated from day 8 to day 10 (p=0.04 q=0.03; Figure 5 B). A significant enrichment was only detected in embryos derived from insemination with fresh spermatozoa (Figure 5). In this stage the TFs E2F-4 8.1×10^−3^, C/EBPgamma 2.57×10^−2^ and ZGPAT 2.69×10^−2^ were significantly enriched, as were the reactome pathways C6 deamination of adenosine, formation of editosomes by ADAR proteins and mRNA editing Ato I conversion (2.39×10^−2^) (Figure 6)

**Figure 5.**
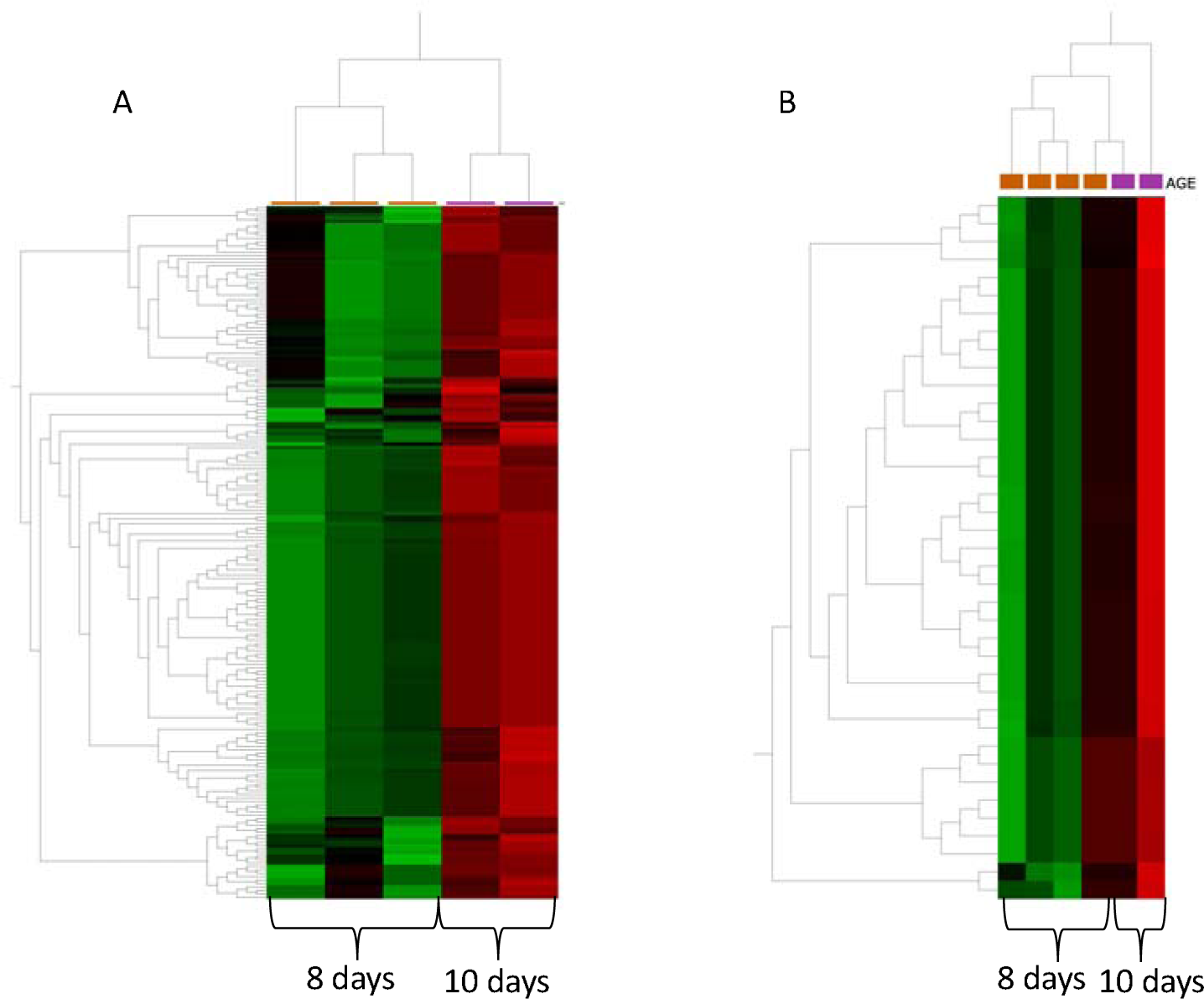
- Heat maps showing the transition from 8 to 10 days of embryo development, A) 202 upregulated expressed transcripts were detected (p=0.04 q=0.1) in embryos obtained after insemination with fresh semen. B) In embryos obtained after insemination with frozen thawed semen only 39 transcripts were upregulated from day 8 to day 10 (p=0.04 q=0.03)

**Figure 6.-.**
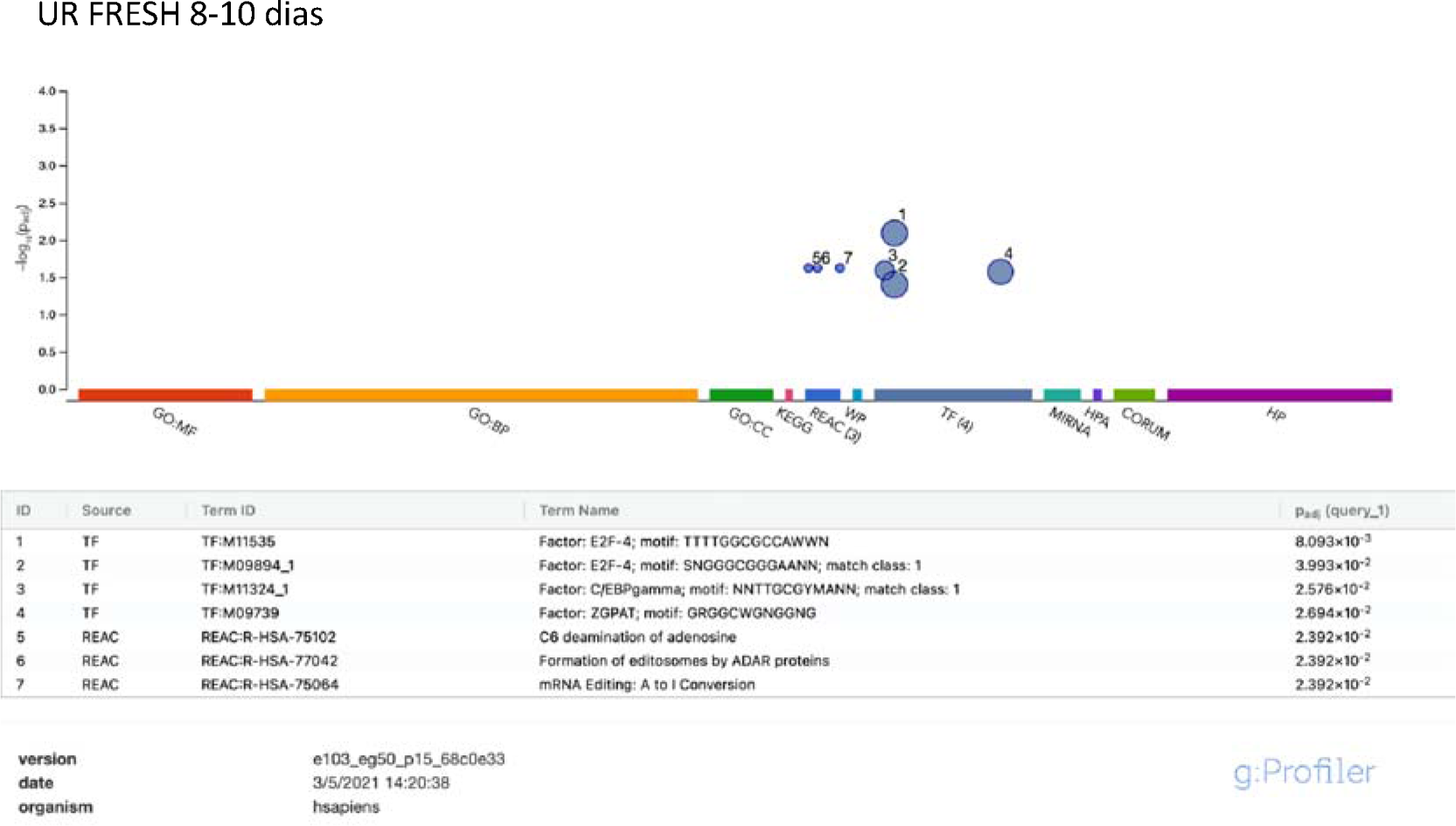
g:GOST multiquery Manhattan plot showing enrichment analysis of the transcripts upregulated of embryos obtained with fresh spermatozoa in the transition from 8 to 10 days after ovulation. Three Reactome pathways and four transcription factors were significantly enriched. The *P* values are depicted on the y axis and in more detail in the results table below present the image.

### Impact of cryopreservation on the development of equine embryos from 10 to 12 days post ovulation

On the other hand, when we compared 10 versus 12 days old embryos it was evident that some genes were differentially expressed, with a group of transcripts upregulated and another downregulated (Fig 7 C). This stage was characterized by a dramatic change in the transcriptome of equine embryos irrespective of the kind of spermatozoa used, either fresh or frozen-thawed.

**Fig 7.-.**
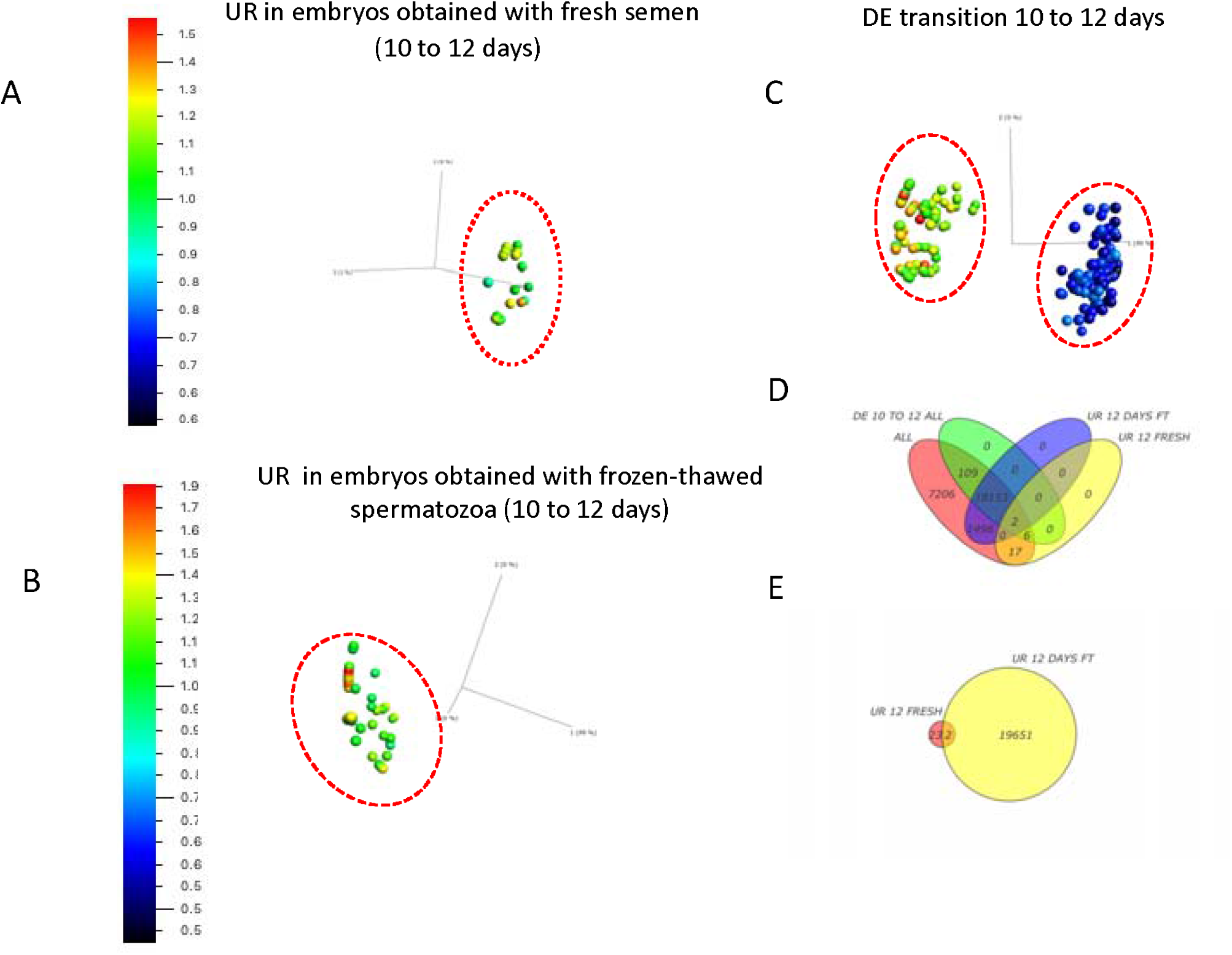
3D Principal component analysis of the changes in the transcriptome in embryos derived from fresh or cryopreserved spermatozoa in the transition from 10 to 12 days of embryo development. A total of 18,270 transcripts were differential expressed in the transition from 10 to 12 days in all embryos (C and D; p=0.02; q=0.03). B) 19651 transcripts were upregulated in embryos resulting from cryopreserved spermatozoa (p=0.04; q=0.07), while only 25 transcripts were upregulated in the transition from 10 to 12 days in embryos obtained with fresh semen (p=0.049; q=0.09; A-D).

A total of 18270 transcripts were differential expressed in the transition from 10 to 12 days in all embryos (Fig 7 C and D; p=0.02; q=0.03); 19651 transcripts were upregulated in embryos resulting from cryopreserved spermatozoa (p=0.04; q=0.07), while, only 25 transcripts were upregulated in the transition from 10 to 12 days in embryos obtained with fresh semen (p=0.049; q=0.09; Figure 7 D-E). Enrichment analysis of these transcripts, after conversion to their human orthologs, revealed dramatic differences between both conditions, with two terms (MIRNAs) enriched in the transition from 10 to 12 days in fresh spermatozoa (has-miR-3150a-3p; 4.2×10^−2^ and has-miR 6763-5p 4.2×10^−2^; Fig 6). On the contrary, embryos derived from cryopreserved spermatozoa showed significant enrichment in many gene ontology terms; 10 for molecular function (MF) and 58 for biological process (BP) (Figure 8 and suppl figures 3-5). The reactome pathway metabolism was significantly enriched (4.48 × 10^−17^), as was the TF TEF-3:EBPbeta (9.48×10^−3^). Detailed description for all the enrichment is provided in supplementary figures. To better disclose the meaning of the increase in the number of annotations observed, we performed PANTHER overrepresentation tests, and found significantly underrepresented annotations in the categories “Protein class” (Table 1) and “Molecular Function” (Table 2). Interestingly this analysis showed many annotations underrepresented, including the protein classes gene-specific transcriptional regulator (PC00264; p=3.52E-03, FDR=3.77E-02), DNA-binding transcription factor (PC00218: p=2.52E-03; FDR=2.52E-03), zinc finger transcription factor (PC00244; p=9.42E-05, FDR=2.60E-03) and chromatin/chromatin-binding, or -regulatory protein (PC00077; p=2.94E-04, FDR=6.31E-03). Among the GO terms (MF) underrepresented in transcripts obtained from embryos obtained with cryopreserved spermatozoa were transmembrane receptor protein kinase activity (GO:0019199; p=2.35E-05, FDR=7.99E-03) and transmembrane receptor protein tyrosine kinase activity (GO:0004714; p=2.08E-05, FDR=7.62E-03).

**Table 1.-.**
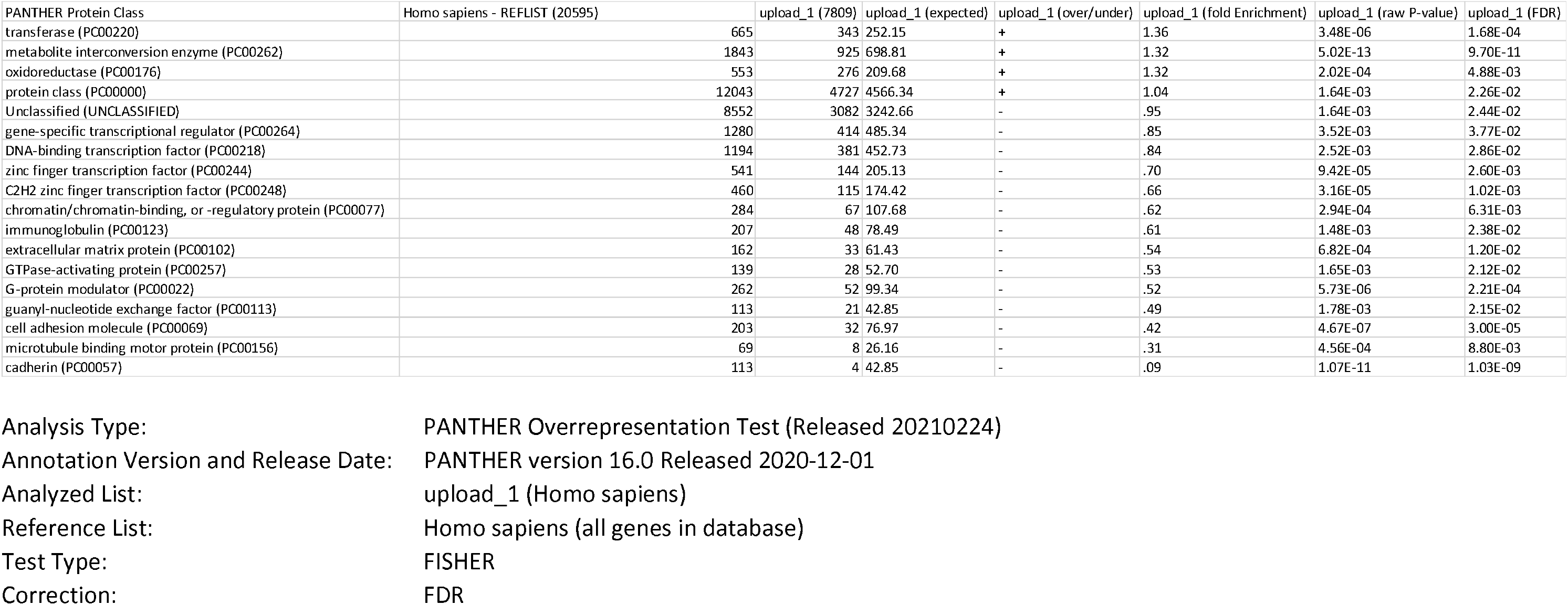
Panther overrepresentation test (Protein class) of transcripts upregulated in the transition from 10 to 12 days in embryos derived from cryopreserved spermatozoa.

**Table 2.** Panther overrepresentation test (Molecular Function) of transcripts upregulated in the transition from 10 to 12 days in embryos derived from cryopreserved spermatozoa.

| GO molecular function complete | Homo sapiens - REFLIST (20595) | upload_1 (7809) | upload_1 (expected) | upload_1 (over/under) | upload_1 (fold Enrichment) | upload_1 (raw P-value) | upload_1 (FDR) |
| --- | --- | --- | --- | --- | --- | --- | --- |
| oxidoreductase activity (GO:0016491) | 759 | 367 | 287.79 | + | 1.28 | 1.17E-04 | 2.95E-02 |
| identical protein binding (GO:0042802) | 2079 | 906 | 788.29 | + | 1.15 | 2.49E-04 | 4.76E-02 |
| protein binding (GO:0005515) | 14300 | 5901 | 5422.13 | + | 1.09 | 9.34E-25 | 1.48E-21 |
| catalytic activity (GO:0003824) | 5778 | 2376 | 2190.84 | + | 1.08 | 8.78E-05 | 2.46E-02 |
| binding (GO:0005488) | 16484 | 6614 | 6250.23 | + | 1.06 | 6.78E-20 | 8.08E-17 |
| molecular_function (GO:0003674) | 18235 | 7265 | 6914.16 | + | 1.05 | 8.53E-31 | 4.07E-27 |
| DNA binding (GO:0003677) | 2483 | 791 | 941.48 | - | .84 | 4.66E-06 | 2.22E-03 |
| adenyl nucleotide binding (GO:0030554) | 1572 | 468 | 596.05 | - | .79 | 1.28E-06 | 7.64E-04 |
| adenyl ribonucleotide binding (GO:0032559) | 1559 | 463 | 591.13 | - | .78 | 1.16E-06 | 7.89E-04 |
| ATP binding (GO:0005524) | 1495 | 437 | 566.86 | - | .77 | 4.52E-07 | 3.59E-04 |
| nucleoside-triphosphatase regulator activity (GO:0060589) | 522 | 132 | 197.93 | - | .67 | 1.64E-05 | 6.52E-03 |
| GTPase regulator activity (GO:0030695) | 478 | 114 | 181.24 | - | .63 | 3.09E-06 | 1.64E-03 |
| Unclassified (UNCLASSIFIED) | 2360 | 544 | 894.84 | - | .61 | 8.53E-31 | 2.03E-27 |
| GTPase activator activity (GO:0005096) | 277 | 60 | 105.03 | - | .57 | 3.65E-05 | 1.09E-02 |
| ATPase activity (GO:0016887) | 488 | 92 | 185.03 | - | .50 | 3.45E-11 | 3.30E-08 |
| helicase activity (GO:0004386) | 159 | 29 | 60.29 | - | .48 | 1.08E-04 | 2.85E-02 |
| calcium ion transmembrane transporter activity (GO:0015085) | 135 | 24 | 51.19 | - | .47 | 2.45E-04 | 4.87E-02 |
| motor activity (GO:0003774) | 131 | 18 | 49.67 | - | .36 | 7.81E-06 | 3.39E-03 |
| microtubule motor activity (GO:0003777) | 69 | 7 | 26.16 | - | .27 | 1.47E-04 | 3.50E-02 |
| transmembrane receptor protein kinase activity (GO:0019199) | 81 | 8 | 30.71 | - | .26 | 2.35E-05 | 7.99E-03 |
| transmembrane receptor protein tyrosine kinase activity (GO:0004714) | 62 | 4 | 23.51 | - | .17 | 2.08E-05 | 7.62E-03 |
| extracellular matrix structural constituent conferring tensile strength (GO:0030020) | 41 | 2 | 15.55 | - | .13 | 2.29E-04 | 4.76E-02 |
| microfilament motor activity (GO:0000146) | 29 | 0 | 11.00 | - | < 0.01 | 1.91E-04 | 4.14E-02 |
| ATP-dependent microtubule motor activity (GO:1990939) | 35 | 0 | 13.27 | - | < 0.01 | 2.58E-05 | 8.22E-03 |
| glutamate receptor activity (GO:0008066) | 28 | 0 | 10.62 | - | < 0.01 | 1.75E-04 | 3.97E-02 |

Analysis Type
PANTHER Overrepresentation Test (Released 20210224)
Annotation Version and Release Date: GO Ontology database DOI: 10.5281/zenodo.4495804 Released 2021-02-01
Analyzed List: upload\_1 (Homo sapiens)
Test Type: FISHER
Correction: FDR

**Fig 8.-.**
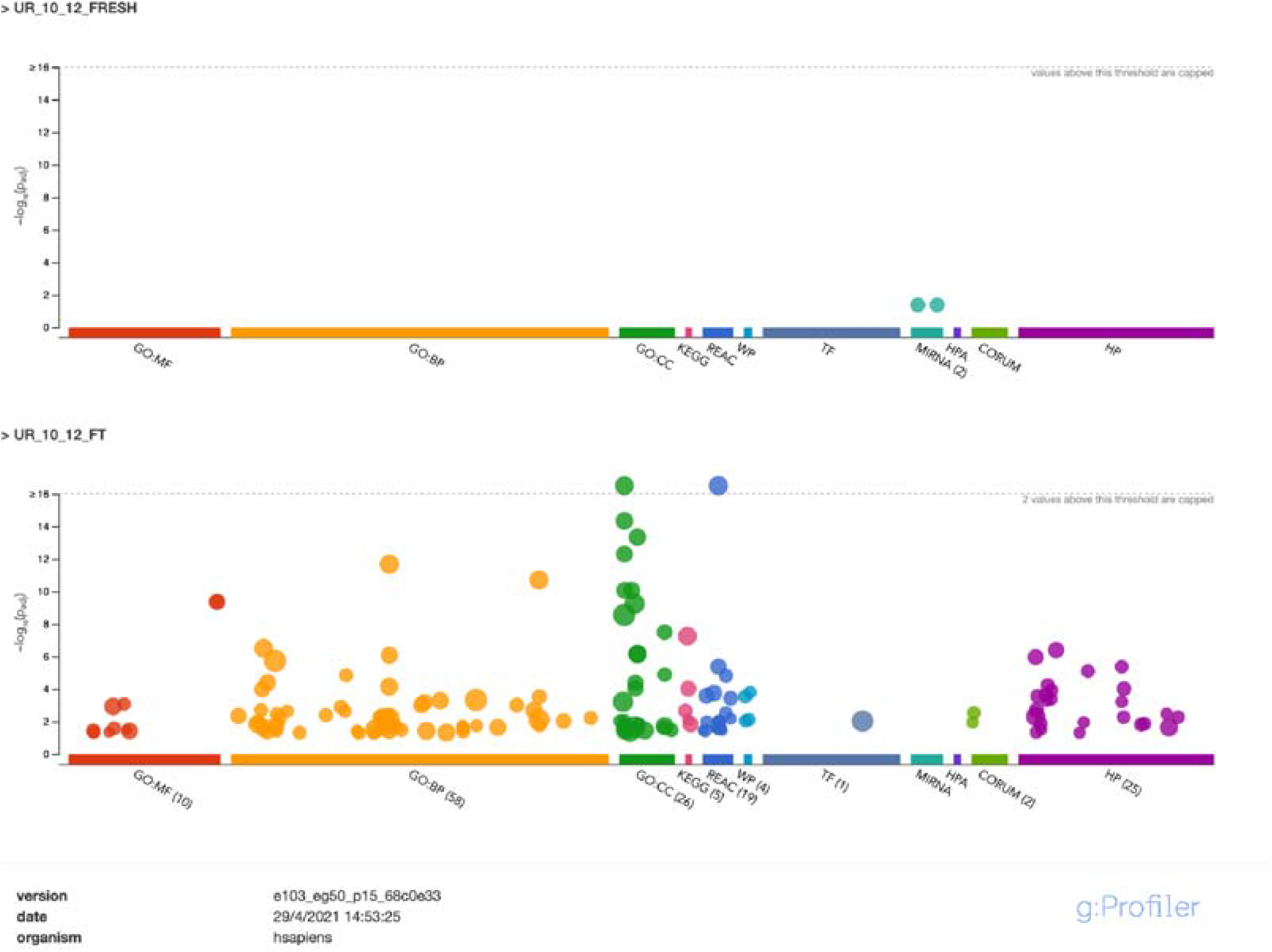
g:GOST multiquery Manhattan plot showing comparative enrichment analysis of equine embryo transcripts under two different experimental conditions; embryos derived from inseminations with fresh or cryopreserved spermatozoa. Gene Ontology terms (GO) for molecular function (MF) are in red, for Biological Process (BP) in orange, and for cellular component (CC) in green. The P values are depicted in the y axis and more detailed in the result table below the image. Enrichment analysis was performed using human orthologs

## DISCUSSION

In the present study we investigated the overall impact of the use of cryopreserved spermatozoa on the transcriptome of equine embryos, and whether the use of frozen-thawed spermatozoa may impact the development of equine embryos at two stages, 8 to 10 days, and 10 to 12 days after ovulation. Cryopreservation is known to cause significant sperm death (Pena et al. 2011; Munoz et al. 2016; Ortega Ferrusola et al. 2017), and the surviving population experiences accelerated senescence characterized by mitochondrial malfunction and redox deregulation leading to reduced lifespan of cryopreserved spermatozoa (Martin Munoz et al. 2015; Ortega-Ferrusola et al. 2019; Ortiz-Rodriguez et al. 2019a; Ortiz-Rodriguez et al. 2021). One important aspect is the role of the spermatozoa on embryo development (Teperek et al. 2016). A recent study of the human sperm proteome (Castillo et al. 2018), identified 103 proteins with known roles in fertilization and 93 with roles in early embryo development. Additionally, 560 sperm proteins were found as involved in modulating gene expression by regulation of transcription, DNA methylation, histone post-translational modifications and non-coding RNA biogenesis (Castillo et al. 2015; Castillo et al. 2018). Some of these proteins may be critical for gene expression regulation after embryo genome activation; the integrative analysis of the sperm, oocyte and blastocyst proteomes and transcriptomes revealed a set of embryo proteins with an exclusive paternal origin, some of which are crucial for correct embryogenesis and, possibly, for modulation of the offspring phenotype (Castillo et al. 2018). In relation to this, cryopreservation is known to impact the sperm proteome (Bogle et al. 2017; Martin-Cano et al. 2020b; Gaitskell-Phillips et al. 2021b; Gaitskell-Phillips et al. 2021a). Taking all these findings in account, is likely that embryos produced with cryopreserved spermatozoa may experience alterations resulting of changes induced by cryopreservation in the proteome of the spermatozoa. Overall, we observed a downregulation of numerous transcripts in embryos resulting from inseminations with cryopreserved spermatozoa in relation to those originated after insemination with fresh spermatozoa. This fact can be easily explained due to the damage induced by cryopreservation to sperm proteins with regulatory roles in embryo development; interestingly and further supporting this hypothesis, all these downregulated transcripts were transcription factors considered essential for embryo development from a very early stage (Godini and Fallahi 2019). Many of the transcription factors found downregulated in embryos obtained using cryopreserved spermatozoa have crucial functions in embryo development, explaining the delayed development and increased embryo mortality attributed to inseminations with cryopreserved semen (Jia et al. 2015). In relation to this, factors found downregulated in embryos derived from insemination with cryopreserved spermatozoa included the Krüppel-like transcription factor3 (KLF)13. This is a member of the Krüppel-like family of transcription factors that controls many growth and developmental processes (Zhou et al. 2007). KLF13 has an important role sensitizing the endometrium to progesterone being crucial for pregnancy initiation and maintenance (Martin et al. 2001; Pabona et al. 2010; Heard et al. 2012; Chen et al. 2015; Grasso et al. 2018). Other members of this family of Kruppel-like factors, including KLF3, KLF14, KLF15 and KLF17 were found downregulated in embryos derived from cryopreserved spermatozoa. Among the discriminant variables detected we observed the proteins the *chromatin-remodeling ATPase INO80*, that has important roles in transcriptional regulation, and DNA replication and repair (Jin et al. 2005; Bakshi et al. 2006; Cai et al. 2007). This gene has a specific regulatory effect on the viability, migration and invasion of trophoblast cells and the knockout mouse results in embryonic lethality, this gene has other important functions in embryonic development (Wang et al. 2014; Qiu et al. 2016; Xian et al. 2021). Another transcript identified was *The mitochondrial mRNA pseudouridine synthase RPUSD3 that catalyzes uridine to pseudouridine isomerization*, that is essential for specific mitochondrial mRNAs translation (*Antonicka et al. 2017*). Downregulation of these proteins argues in favor of a compromise in the development of embryos obtained with cryopreserved spermatozoa; further supporting this hypothesis the visualization of integration networks using the Cytoscape app Clue go revealed annotations like regulation of nuclear division, mitotic DNA damage checkpoint and positive regulation of molecular mediator of immune response significantly affected by cryopreservation.

When we investigated whether embryo development was affected when cryopreserved spermatozoa was used, we found a dramatic difference in the number of transcripts upregulated in embryos derived from fresh spermatozoa, respect embryos derived from inseminations with cryopreserved spermatozoa in the transition from 8 to 10 days, this fact may help to explain why embryos obtained with cryopreserved spermatozoa experience a delay in development in this particular interval (Stout 2006). However, from 10 to 12 days, a higher number of transcripts were upregulated in embryos obtained with cryopreserved spermatozoa. These data may indicate that embryos derived from cryopreserved spermatozoa may lose control of transcription control, as has been described in low quality human embryos (Groff et al. 2019). One of the possible explanations for this finding may be delayed development of these embryos, representing late activation of transcripts. While the enrichment of transcripts related to metabolism may argue in favor of this argument, the GO terms involved in post transcriptional gene silencing were highly enriched. Moreover, the human phenotype ontology (https://hpo.jax.org/app/) terms were enriched in embryos obtained with cryopreserved spermatozoa, including terms related to abnormal metabolism, which may suggest an abnormal gene activation in these embryos. To further explore the biological meaning of these findings we also conducted Panther overrepresentation tests, to found a significant underrepresentation of protein classes involved in transcriptional regulation, and different transcription factors. Moreover, underrepresentations of protein classes related to immune functions were also observed which argues in favor of abnormal gene activation, reinforced by the molecular function GO terms related to cellular signaling like receptor protein kinase and protein tyrosine kinase activities highly underrepresented.

In sum, cryopreservation seriously influences the transcriptome of early embryos, having a direct impact in all embryos derived from cryopreserved spermatozoa, characterized by downregulation of numerous transcription factors. In addition, cryopreservation also impacts embryo development in the transition from 8 to 10 days, and from 10 to 12 days. Overall, our findings provide strong evidence that insemination with cryopreserved spermatozoa may compromise early embryo development, probable due to cryo-induced modifications in sperm proteins. Identification of which specific factors may cause such changes in the sperm proteome induced by cryopreservation is expected.

## ACKNOLEDGEMETS

The authors received financial support for their investigations from the Ministerio de Ciencia-FEDER, Madrid, Spain, grant <u>AGL2017-83149-R and PID2019-107797RA-I00/AEI/10.13039/501100011033,</u> Junta de Extremadura-FEDER (GR18008), JMOR holds a Predoctoral grant from Junta de Extremadura-FEDER (PD 18005) GGPH holds a PhD grant from the Ministry of Science, Madrid, Spain (PRE 2018-083354).

